# Role for Ribosome-Associated Complex and Stress-Seventy subfamily B (RAC-Ssb) in integral membrane protein translation

**DOI:** 10.1101/179564

**Authors:** Ligia Acosta-Sampson, Kristina Döring, Yuping Lin, Vivian Y. Yu, Bernd Bukau, Günter Kramer, Jamie H. D. Cate

## Abstract

Targeting of most integral membrane proteins to the endoplasmic reticulum is controlled by the signal recognition particle (SRP), which recognizes a hydrophobic signal sequence near the protein N-terminus. Proper folding of these proteins is monitored by the unfolded protein response, and involves protein degradation pathways to ensure quality control. Here, we identify a new pathway for quality control of major facilitator superfamily transporters that occurs before the first transmembrane helix–the signal sequence recognized by SRP–is made by the ribosome. Increased rates of translation elongation of the N-terminal sequence of these integral membrane proteins can divert the nascent protein chains to the ribosome-associated complex (RAC) and Stress-Seventy Subfamily B (Ssb) chaperones. We also show that quality control of integral membrane proteins by RAC-Ssb couples translation rate to the unfolded protein response, which has implications for understanding mechanisms underlying human disease and protein production in biotechnology.

---

Integral membrane protein open reading frames (ORFs) make up approximately 20-30% of eukaryotic genomes (1). Membrane protein (MP) biogenesis is a highly coordinated, multi-phase process in which many of the steps are thought to be kinetically controlled. Early in membrane protein synthesis, the ribosome nascent chain (MP-RNC) complex is identified by the signal recognition particle (SRP) and targeted to the endoplasmic reticulum (ER), for insertion of the MP into the ER membrane through the Sec translocon (2,3). For membrane proteins, exposure of the first transmembrane helix (TM1) at the ribosome peptide exit site serves as the signal for SRP binding, though SRP can be recruited to the ribosome while TM1 is still inside the ribosome exit tunnel (4-8). SRP-target binding is also mediated by the nascent-polypeptide associated complex (NAC), which prevents non-specific binding of SRP to non-secretory substrates, and deters antagonistic interactions between SRP and N-terminal-modifying enzymes during scanning of nascent polypeptides (9-11). These early targeting steps are crucial for proper membrane protein insertion into the ER and to prevent proteostatic stress due to the aggregation of the highly hydrophobic transmembrane helices of membrane proteins in the cytosol (5).

The contributions of kinetics in controlling nascent chain targeting has remained unclear. SRP-mediated attenuation of translation elongation has been proposed to extend the time window for SRP targeting in eukaryotes and is necessary for efficient translocation in human cells (12-14). However, a recent study using SRP-mediated selective ribosome profiling (SeRP) in yeast did not find evidence for general SRP-mediated pausing on secretory nascent chain substrates (7). Instead, computational analyses of mRNAs encoding membrane proteins and secreted proteins have identified clusters of rare codons that modulate their translation elongation rates to facilitate constructive interactions with SRP (15,16). Tuller and colleagues found that transcripts from several Gene Ontology (GO) categories, including those composed of secreted and membrane proteins, had a 5’-end slow translational ramp of 30-50 codons comprised of non-optimal codons that optimizes ribosome spacing and reduces the chances of ribosomal ‘traffic jams’ (17). Moreover, a genome-based analysis of N-terminal signal-sequences in Sec-dependent secreted proteins from *E. coli* showed that these sequences had a high frequency of non-optimal codons (18). Downstream from the N-terminus, Pechmann and colleagues showed through *in silico* sequence analysis and ribosome profiling data that secreted and membrane proteins had REST (mRNA-encoded slowdown of translation) elements of non-optimal codons 30-40 codons downstream of the signal-sequence/TM1 that assisted SRP interactions through slowing translation elongation (16). All of these studies relied on computational methods to discern patterns of codon usage bias. The earliest metric designed to detect this bias was the Codon Adaptation Index (CAI), which is based on the codon usage frequencies of highly expressed genes in a given species (19). The CAI adequately predicts gene expression levels in *E. coli* and *Saccharomyces cerevisiae,* whose codon usage frequencies are positively correlated with isoaccepting tRNA abundances (20). Subsequently, a wide-ranging scale, the tRNA Adaptation Index (tAI), was introduced that better described the relationship between tRNA pools and codon usage patterns for unicellular and multicellular organisms. The tAI integrates the tRNA gene copy number and Crick’s wobble rules for codon-anticodon pairing (21). Global tAI has a strong correlation with high expression levels for cytosolic proteins, but this correlation is weak for membrane and secretory proteins in *S. cerevisiae* (15).

At the protein level, codon usage preferences also play an important role in regulating membrane protein function, folding and structural stability. Several pathogenic synonymous polymorphisms in the human *MDR1* gene, which encodes the P-glycoprotein drug efflux transporter, have been identified that alter the structure and efflux activity of the transporter (22,23). Misfolding and aggregation of secreted and membrane proteins trigger the ER quality control system characterized by two separate processes, the unfolded protein response (UPR), a signaling pathway that monitors ER homeostasis and responds to problems during membrane protein biogenesis at the ER, and the ER-associated degradation pathway (ERAD), which targets misfolded proteins in the ER lumen and membrane for proteasomal degradation (24-26).

Among the different classes of MPs, the major facilitator superfamily (MFS) is a ubiquitous family of secondary transporters that employ electrochemical gradients to drive substrate translocation through the membrane [Transporter Classification Database (TCDB) #2.A.1; (27)] (28). It is the largest family of secondary transporters including up to 25% of membrane transporters in prokaryotes (29). Most members range in size from 400 to 600 amino acids and are organized into 12 transmembrane α-helices. MFS transporters are important drug and therapeutic targets because they transport a variety of substrates including simple carbohydrates, peptides, oligosaccharides and lipids (30). These transporters can be divided based on their use of three distinct transport mechanisms: uniporters, symporters and antiporters (28). The sugar porter subfamily is one of the largest and most widely studied subfamilies of MFS transporters, and includes the human *SCLC2* (GLUT) family of membrane transporters, the *HXT* family of hexose transporters from *S. cerevisiae,* and the oligosaccharide family of transporters in *Neurospora crassa.* The transcriptional regulation of these transporters has been extensively studied in relation to extracellular substrate concentrations and tissue specificities. However codon preferences in mRNAs encoding these transporters, and regulation of their proper targeting to the ER have not been studied to date (31-34). Here we present experimental evidence for a new quality control mechanism that couples translation rates of the N-terminus of MFS transporters to the chaperone machinery at the ribosome exit tunnel, which may contribute to proper SRP interactions with these ribosome nascent chains.

## RESULTS

### Codon-optimization of the N. crassa CDT-1 transporter

The production of biofuels from plant cell walls requires the breakdown of cellulose into soluble glucose. However the cellulase enzymes that cleave cellulose are inhibited by the product cellobiose, a β-1,4-linked dimer of glucose (35). We previously showed that heterologous co-expression of the *Neurospora crassa* CDT-1 MFS transporter and intracellular β-glucosidase GH1-1 (NCU00801 and NCU00130, respectively) enabled *Saccharomyces cerevisiae* cells to consume cellobiose directly, which could circumvent cellulase inhibition during biofuel production (36) (Figure 1A). However, the CDT-1 transporter is slow relative to glucose transporters in yeast, and remains a bottleneck in the cellobiose consumption pathway (37). To improve cellobiose consumption rates, we compared codon usage bias in the endogenous *cdt-1* sequence derived from *N. crassa* versus either the *N. crassa* genomic background or the *S. cerevisiae* genomic background. Using the %MinMax algorithm (38), which uses a sliding window to compare a coding sequence to the most common (%Max) or the most rare codons (%Min) from the genomic codon usage tables of a particular organism, revealed that the *Neurospora* coding sequence is optimal in *N. crassa*, but sub-optimal in *S. cerevisiae,* with most of the open reading frame composed of codons used less frequently in yeast (Figure 1B). Other codon usage metrics such as the codon adaptation index (CAI) and the tRNA adaptation index (tAI) yielded similar results (Figure S1)(19,21,39).

**FIGURE 1.**
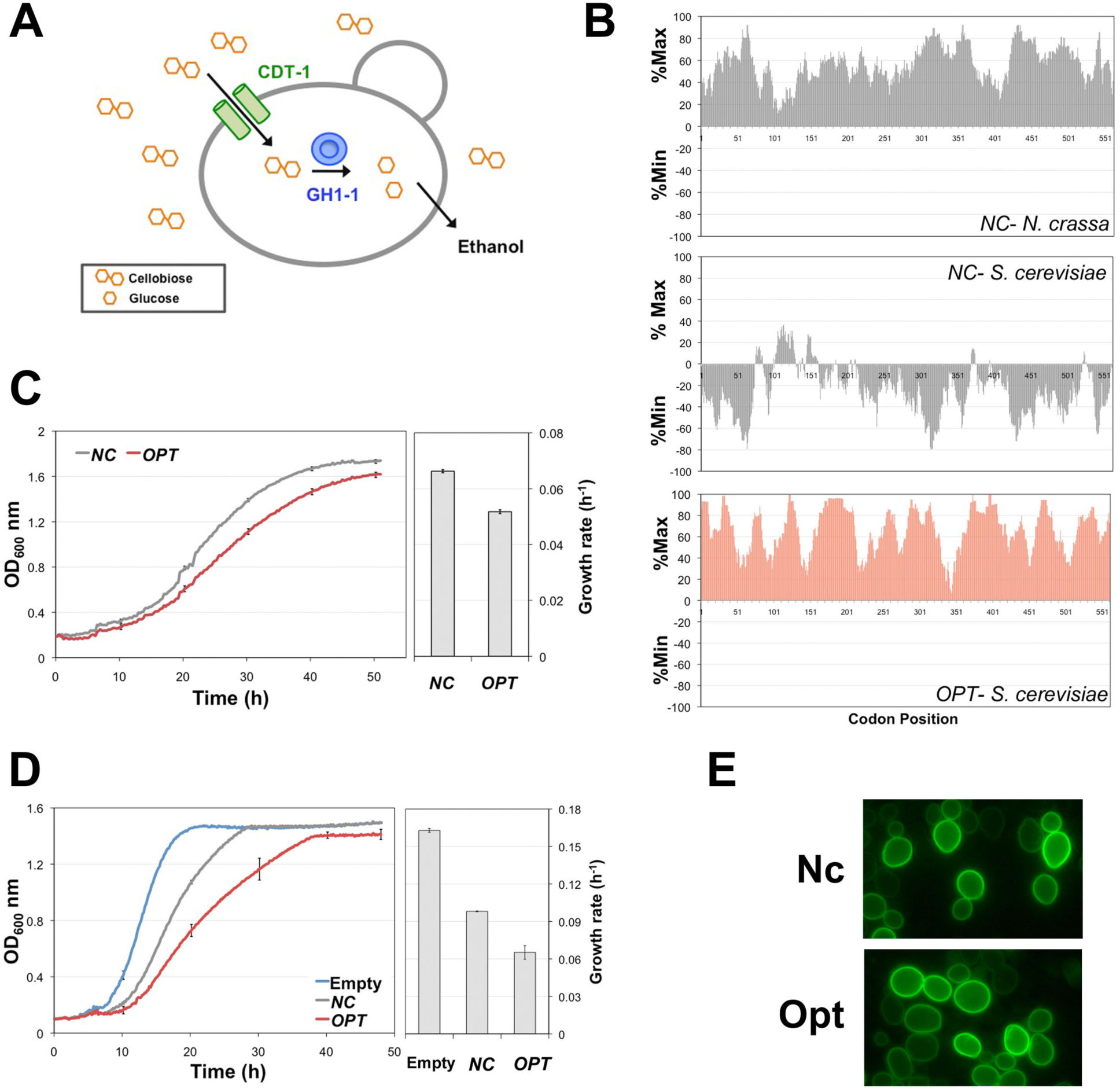
Codon-optimized CDT-1 and growth phenotypes. (*A*) Schematic of the cellobiose-utilization pathway in *S. cerevisiae*. The CDT-1 transporter imports cellobiose into the cell and the GH1-1 β-glucosidase hydrolyzes cellobiose into two glucose monomers, which then enter the glycolytic pathway. (*B*) %MinMax profiles for *NC* using *S. cerevisiae* and *N. crassa* codon usage tables and %MinMax profile for *OPT* using the *S. cerevisiae* codon usage table. (*C*) Growth curves and growth rates for D452-2 cells expressing *NC* or *OPT* and *GH1-1* in media containing cellobiose as the sole carbon source, under aerobic conditions. Growth rates were calculated from the exponential growth phase and were 0.07 h^−1^ for cells expressing *NC* and 0.05 h^−1^ for cells expressing *OPT*. (*D*) Growth curves and growth rates for D452-2 cells expressing *NC* or *OPT* in glucose-containing media. Growth rates were: 0.16 h^−1^ for cells transformed with empty plasmid; 0.1 h^−1^ for cells with pRS316-*NC*; and 0.065 h^−1^ for cells with pRS316-*OPT*. All growth rates shown represent the mean growth rate from three biological replicates, with the corresponding standard error. (*E*) Cellular localization of Nc and Opt. Epi-fluorescent microscope images of *S. cerevisiae* strain D452-2 cells with Nc-GFP (Nc) or Opt-GFP (Opt) transporters.

We therefore optimized the codons in the *cdt-1* sequence to better match the codon preferences of *S. cerevisiae*. After codon optimization of *cdt-1*, the CAI metric showed that the optimized version consists of the most common codons, whereas the *Neurospora*-derived sequence had much lower codon preferences (Figure S1A) (19,39). The tRNA adaptation index tAI was also higher for the optimized *cdt-1* sequence (Figure S1B)(21). Notably, improvement in the CAI and tAI still resulted in sequence identity at the nucleotide level for the *Neurospora*-derived and codon-optimized open reading frames of 76.2 % (Clustal Omega alignment, Figure S2).

### Codon-optimized CDT-1 causes a slow-growth phenotype

We previously found that codon-optimization of *GH1-1* improved fermentation rates under aerobic and anaerobic conditions (40). However, we found that the codon-optimized version of the CDT-1 transporter (here denoted *OPT* for the mRNA and Opt for the expressed transporter) resulted in reduced cellobiose uptake and slower growth in cellobiose by *S. cerevisiae* relative to the transporter with *Neurospora*-derived coding sequence (hereafter *NC* for the mRNA, Nc for the expressed transporter) (Figure 1C, Figure S3). Additionally, in the course of preparing cultures for cellobiose growth assays, we noticed that single-colony seed cultures of *S. cerevisiae* cells constitutively expressing *OPT* grew significantly slower in glucose media than did their counterparts expressing *NC*, even though both genes were expressed using the same vector with identical promoter and terminator (pRS316-*P*_*THD3*_-*CYC1tt*), which also result in identical 5’- and 3’-untranslated regions (UTRs). We therefore set up a growth assay in optimized Minimal Media plus glucose (oMM-Ura), which revealed that yeast expressing *OPT* grew 0.6-times the rate of cells expressing *NC*, and 0.3-times the rate of cells carrying only the empty vector (Figure 1D). Given these results, we suspected that codon-optimizing *cdt-1* led to improper localization of the transporter or to protein aggregation. However, fluorescent microscopy of cells expressing either Opt or Nc transporter with eGFP fused to the C-terminus revealed identical localization patterns for both proteins. The GFP fluorescence signal was present predominantly at the plasma membrane with very little to no intracellular fluorescence (Figure 1E).

We wondered whether the transporter expressed from *OPT* was misfolded, but not severely enough to trigger ER-associated degradation (ERAD) and prevent trafficking to the plasma membrane. For example, *MDR1* mutants with synonymous polymorphisms that led to conformational changes and impaired drug-efflux activity in the P-glycoprotein transporter had proper plasma membrane localization (23). The presence of a partially misfolded population of Opt could explain the lower cellobiose uptake rates and slower growth in cellobiose (Figure 1C, Figure S3). To test for more subtle conformational destabilization of the CDT-1 transporter, we compared the thermal stability of transporter encoded by *NC* or *OPT* via fluorescence size-exclusion chromatography (FSEC-TM) (41). In FSEC-TM, eGFP is fused to the C-terminus of the target membrane protein in order to use GFP fluorescence as a proxy for absorbance at 280 nm during size-exclusion chromatography. Importantly, while the peak heights for Nc–GFP or Opt–GFP decreased with temperature, the height of the GFP-only peak decreased marginally, indicating that GFP did not unfold during these experiments (41)(Figure 2A). Plotting the experiments as a thermal melt curve (41) revealed the Tm value for both Nc and Opt transporters to be ∼38 °C (Figure 2B). These results indicate that codon-optimization did not affect the stability of the CDT-1 transporter significantly.

**FIGURE 2.**
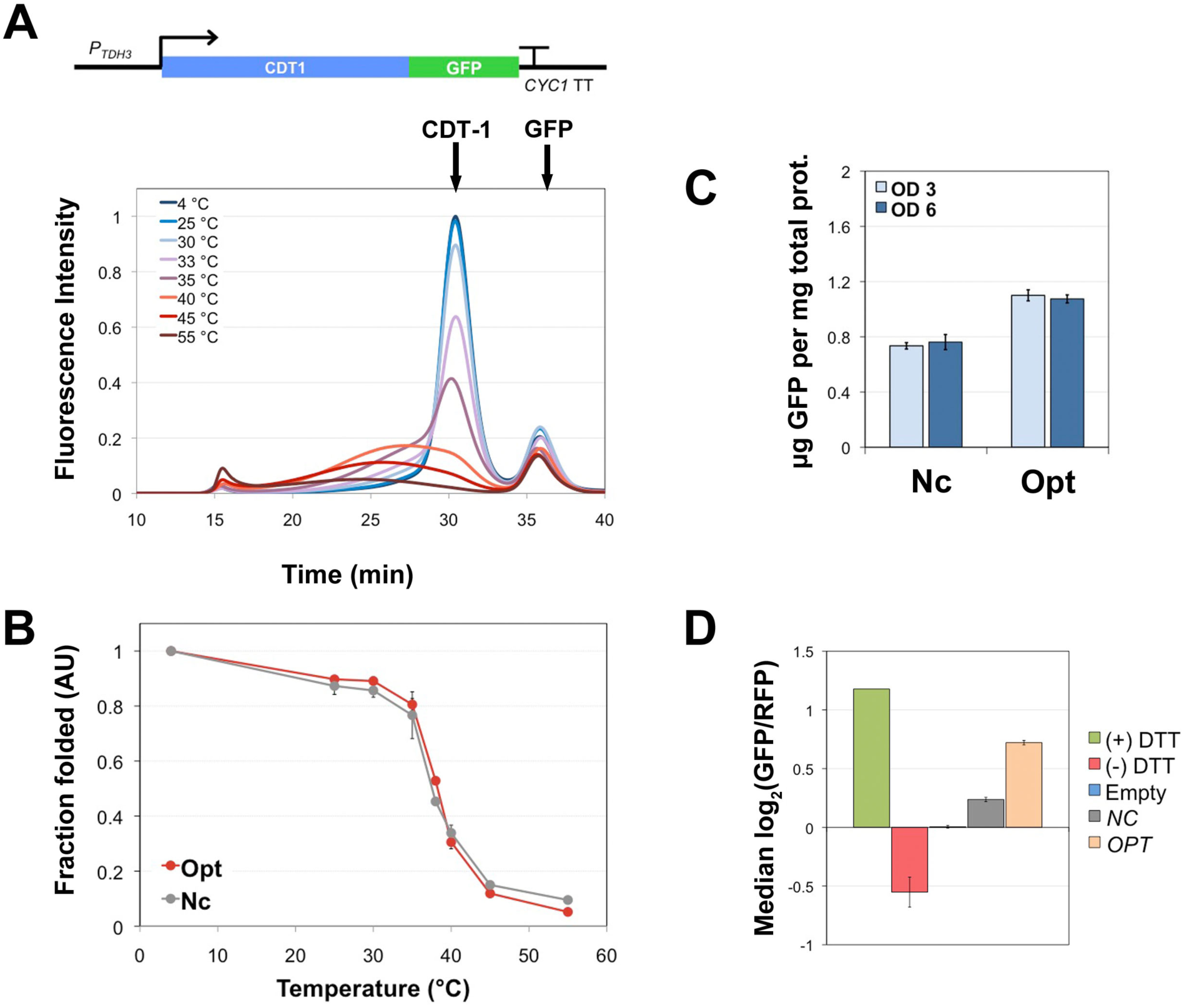
Synonymous codon changes effects on protein stability and the Unfolded Protein Response. (*A*) Measured changes in the thermal stability of Nc via Tm-FSEC. For each temperature tested, a 4 °C incubation control was included. The first eluted peak corresponds to soluble aggregates of Nc-GFP in the void volume of the column. The second eluted peak corresponds to monomeric Nc-GFP and the last eluting peak is free GFP. (*B*) Thermal melting curves for Nc and Opt. The fraction of folded transporter was calculated as the ratio of the peak area for the monomeric Nc/Opt–GFP peak at a given temperature to the peak area of the same monomeric peak for the control (4 °C) within the same elution range to yield a thermal melt curve. The calculated Tm values were 37.8 ± 0.16 °C for Nc, and 38.2 ± 0.14 °C for Opt, with R^2^ values of 0.97 and 0.98, respectively. (*C*) Nc-GFP and Opt-GFP protein concentration measured via GFP fluorescence. Mean values and the corresponding standard errors are shown for three biological replicates. *(D*) *NC-* or *OPT*-induced UPR activation in the D452-2 UPR reporter strain. UPR activation is shown as the median of the log_2_ GFP/RFP ratio. Each bar represents the mean from 4 biological replicates with the corresponding standard error. The ‘Empty’ designation refers to cells transformed with empty plasmid as a control. The baseline for minimal activation of the UPR response was determined by using just the D452-UPR cells to measure no UPR activation [(-) DTT]. The baseline for maximal UPR activation used D452-UPR cells incubated in media containing 5 mM DTT (UPR inducer) for 2-3 h in order to fully induce the UPR [(+) DTT].

We also tested whether codon-optimization of *cdt-1* may have resulted in substantially increased transporter protein expression levels that would tax cell resources, resulting in stress and slower growth. We measured Nc–GFP or Opt–GFP protein concentration in whole cell lysates using GFP fluorescence and found Opt protein levels were 1.5-fold higher than Nc levels (Figure 2C). Protein concentrations for both transporters did not change between mid- and late-exponential phases (OD_600_ = 3 vs. OD_600_ = 6, Figure 2C).

### Codon-optimized CDT-1 activates the unfolded protein response

To assess whether a specific proteostatic stress response pathway was associated with the slow-growth phenotype of cells expressing *OPT*, we examined the ubiquitous unfolded protein response (UPR) (24). We constructed a UPR reporter strain using the yeast background strain used for cellobiose fermentations (D452-UPR), based on the single-cell fluorescence assay developed in (42). Briefly, a bi-partite UPR reporter cassette with GFP is expressed under the control of UPR elements (UPRe) and RFP is expressed constitutively. The levels of GFP fluorescence reveal the level of UPR activation and the levels of RFP fluorescence serve as internal controls for cell size and translation activity to normalize for cell-to-cell variation. UPR activation is represented as the median of log_2_ GFP/RFP ratio for the entire population of cells sampled during flow cytometry.

We transformed plasmids pRS316-*NC* and pRS316-*OPT*, along with an empty pRS316 control plasmid, into strain D452-UPR. We then grew cultures to the same final OD_600_ of 3 (mid-exponential phase) and measured GFP and RFP levels using flow cytometry. Compared to controls with D542-UPR alone, or treated with DTT to fully induce the UPR (42), both *NC* and *OPT* expression activated the UPR response, but the level of activation was significantly higher for *OPT* (Figure 2D). UPR activation has been observed during expression of membrane protein mutants that destabilize the native structure or during over-production of specific membrane proteins for crystallographic purposes (43,44). Although Opt was not misfolded (Figure 2B) and its concentration was not significantly higher than NC (Figure 2C), activation of the UPR stress response suggests that post-transcriptional or translational events during expression of *OPT* may be responsible for the slow-growth phenotype

### The first 30 optimized codons of CDT-1 are responsible for the slow-growth phenotype

Nascent membrane protein insertion and folding in the ER membrane, as well as subsequent steps, such as transmembrane helix (TM) insertion by the Sec61 complex, is under kinetic control (45,46). Thus, synonymous codon changes that could lead to variation in translation elongation speed could affect this highly synchronized process (16,47). SRP mis-recognition or TM mis-insertion could lead to activation of quality control pathways that result in nascent chain degradation and cellular stress (5). We therefore dissected the region of the *OPT* sequence responsible for the slow-growth phenotype, to help pinpoint the step in membrane protein biogenesis that was affected by codon optimization.

We exchanged coding regions from the *NC* sequence into the corresponding regions in *OPT*, using the topology of MFS transporters–which have a basic structural unit consisting of four triple-helix bundles (48)–as a guide. We divided the sequence into four regions, each encompassing one of these triple-helix motifs (Figure 3A), using the Phobius TMH predictor to demarcate the sequence boundaries for each triple-helix bundle (Table S1)(49). We constructed four chimeras in which we sequentially switched the *NC* sequence for each triple-helix bundle into the *OPT* mRNA and used growth assays in glucose to detect if switching any of these regions would rescue the slow-growth phenotype observed in cells expressing *OPT*. We first confirmed that all four transporters expressed from the chimeric mRNAs localized to the plasma membrane (Table S1). Remarkably, switching the first triple-helix bundle sequence in *OPT* for the *NC* counterpart entirely rescued the slow-growth phenotype (Figure 3B). We next switched secondary structural elements individually in the first three-helix bundle region, to yield 7 different chimeras: one for the N-terminal tail region, 3 for the TMs, and 3 for the loops connecting TMs (Table S1). Only the N-terminal tail chimera (*NC* codons 1-72 fused to *OPT* codons 73-579) showed the same growth rate as cells expressing *NC* (Figure 3C).

**FIGURE 3.**
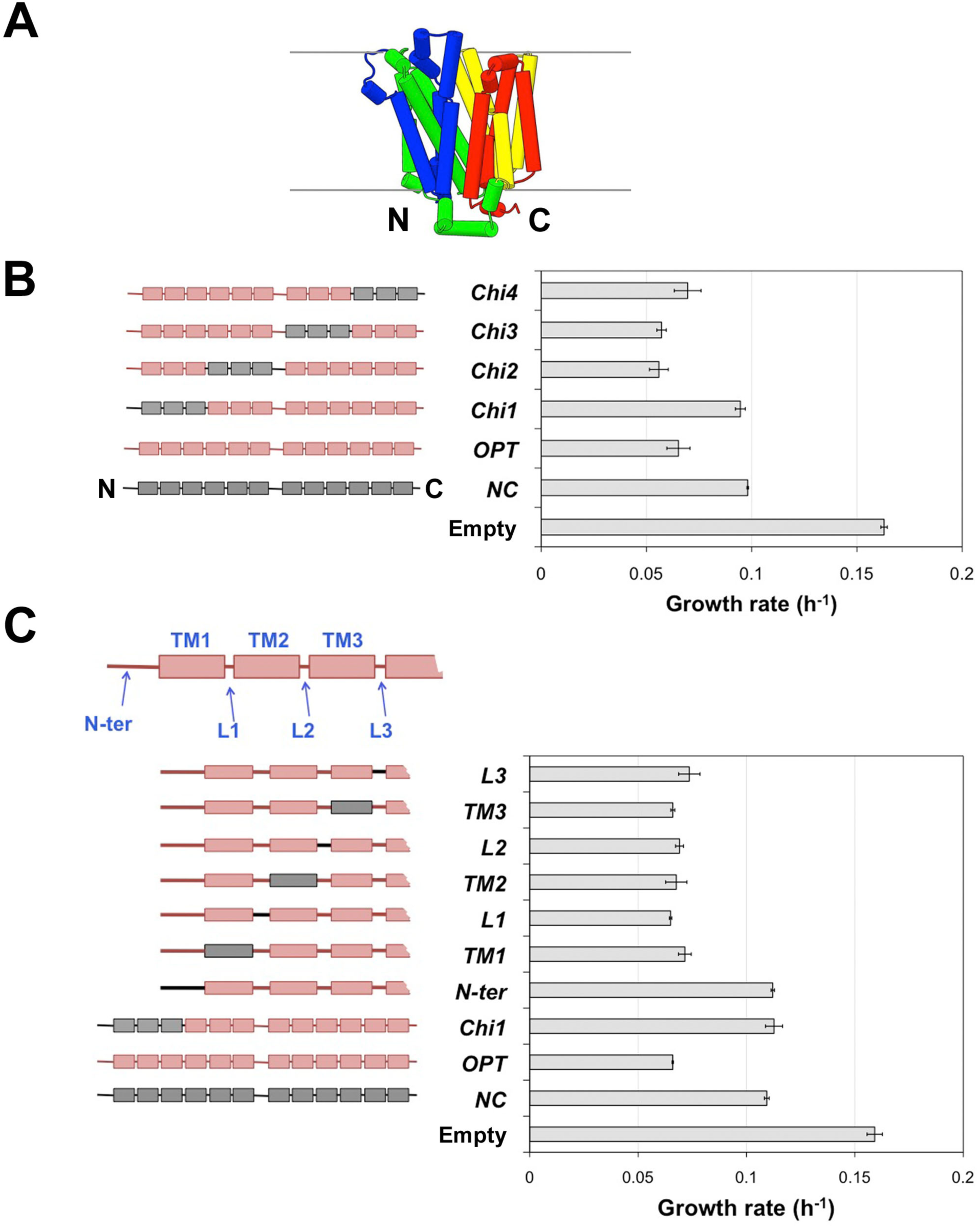
*NC*-*OPT* chimeras. (*A*) Structure of the MFS transporter XylE from *E. coli* [PDB ID: 4GBZ, (90)] showing the four triple-helix motifs. Motifs are colored in succession from the N- to the C-terminus: the first motif is blue; the second is green; the third is yellow; and the fourth is red. (*B*) Growth rates for D452-2 cells expressing triple-helix chimeras. The corresponding region for each triple helix from the *NC* sequence was swapped into the *OPT* sequence. (*C*) Growth rates for D452-2 cells expressing chimeras within the first triple-helix bundle. The first triple-helix motif was divided into individual secondary structural elements and the *NC* sequence was swapped into the corresponding sequence in the *OPT* ORF. All growth rate measurements represent the mean growth rate and standard error from biological triplicates.

We further divided the first 72 codons of *OPT* into 30-codon overlapping windows and swapped in the corresponding *NC* sequences to make three chimeras: the region 1 chimera had *NC* codons 1-30; the region 2 chimera had *NC* codons 21-50; and the region 3 chimera had *NC* codons 41-72 (Figure 4A). Growth assays in glucose showed that only cells expressing the region 1 chimera– with *NC* sequence at the extreme N-terminus–had the same growth rate as cells expressing the intact *NC* sequence (Figure 4A). To further test that the extreme N-terminal sequence of *OPT* was responsible for the slow-growth phenotype, we carried out a reciprocal swap, switching the first 30 codons of the *NC* sequence for the coding sequence of *OPT*. The reciprocal swap of the *OPT* N-terminal coding sequence onto the remaining coding sequence of *NC* conferred the same slow growth phenotype as *OPT* on cells expressing this chimera, as anticipated (Figure 4B). The slow-growth phenotype of cells expressing *OPT* and the region 1 *OPT*/*NC* chimera also correlates with UPR activation in cells expressing these mRNAs (Figure 5A). Thus, the first 30 codons–and possibly the first 20 unique to the region 1 chimeras–were sufficient to establish the slow-growth phenotype and UPR activation induced by codon optimization of CDT-1.

**FIGURE 4.**
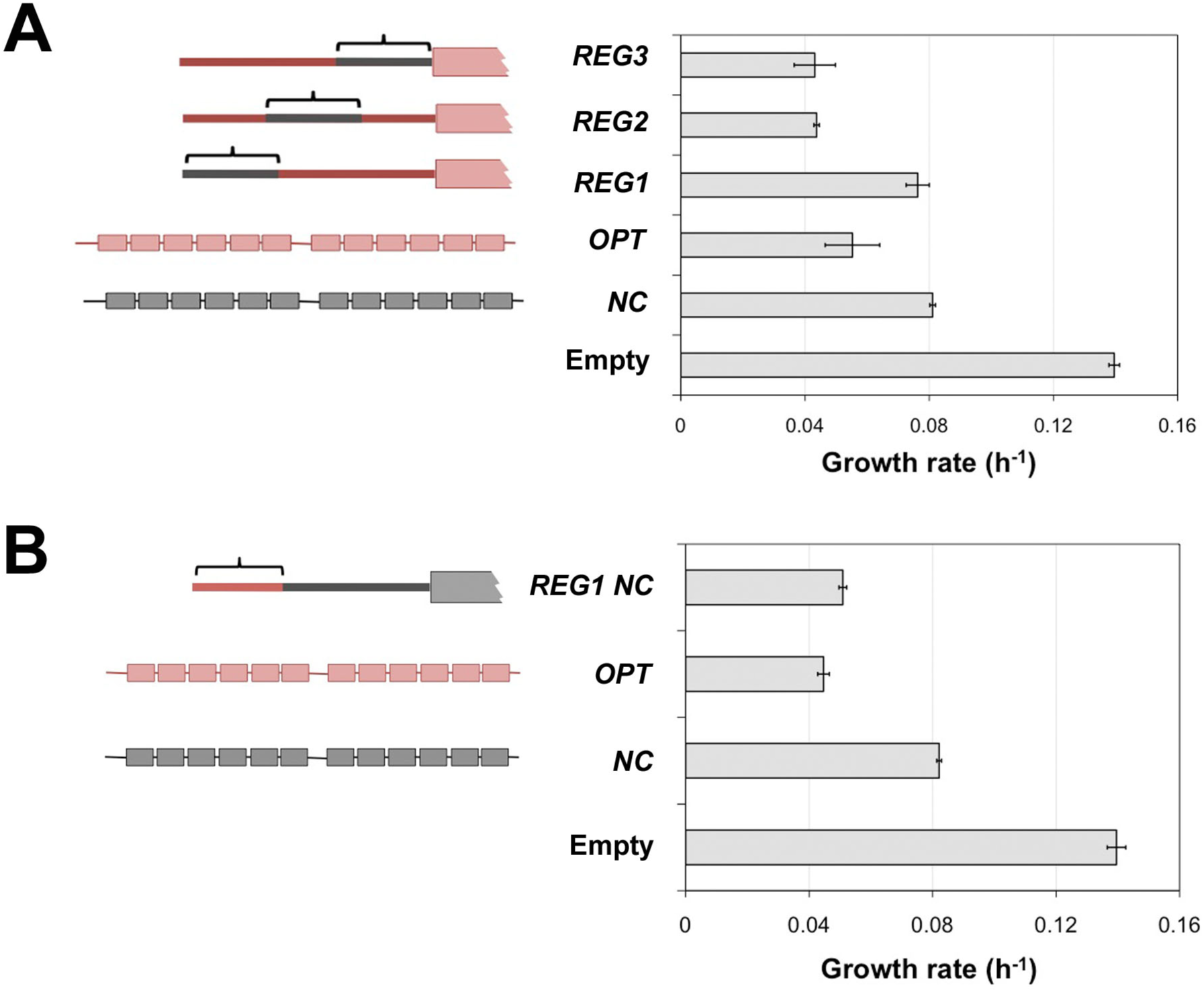
Impact of the N-terminal codons of *OPT* on growth phenotypes. (*A*) Growth rates of cells expressing chimeras within the first 72 codons. The first 72 codons of *OPT* were divided into 30-codon overlapping windows which were replaced with the corresponding *NC* sequences. *REG1* corresponds to codons 1-30 from *NC*, *REG2* to *NC* codons 21-50, and *REG3* to *NC* codons 41-72 (Black brackets). (*B*) Growth rate of cells expressing *OPT* codons 1-30 replacing the *NC* coding sequence (*REG1 NC*). Mean growth rates and standard errors from biological triplicates are shown.

**FIGURE 5.**
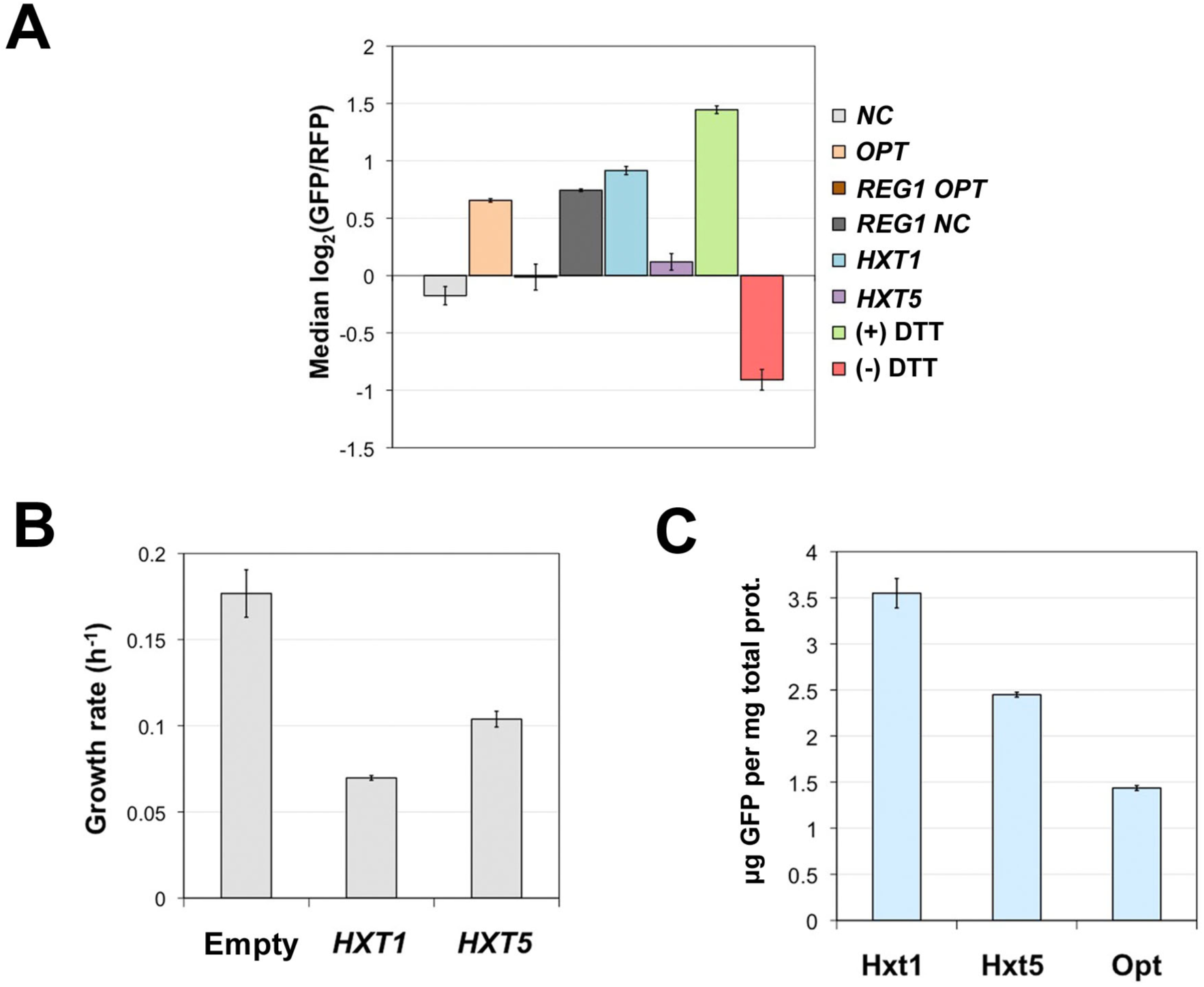
UPR activation in transporters containing the first 30 codons of *OPT*. (*A*) UPR activation levels in cells expressing *NC*, *OPT*, and the *REG1 OPT* and *REG1 NC* chimeras. The level of UPR activation is higher in the presence of region 1 (codons 1-30) from the *OPT* sequence (*REG1 NC*) compared to region 1 of *NC* fused to the remainder of *OPT* (*REG1 OPT*). Mean levels are shown for five biological replicates, with corresponding standard errors. (*B*) Growth rates of BY4741 cells expressing *HXT1* and *HXT5* from plasmids. Growth rates represent the mean growth rate and standard error from three biological replicates. (*C*) Protein concentration of plasmid-expressed Hxt1, Hxt5 and Opt. The Hxt1 levels were 3.55 ± 0.16, Hxt5 levels were 2.45 ± 0.03 and Opt levels were 1.47 ± 0.04 μg GFP/mg total protein. *HXT1* and *HXT5* were expressed from the same expression cassette as *NC* and *OPT*.

To test whether similar high-level expression of other transporters would induce the same growth and UPR effects, we chose two endogenous hexose transporters from *S. cerevisiae*, *HXT1* and *HXT5,* that like CDT-1 are members of the MFS sugar porter subfamily (TCDB #2.A.1.1), and that are not aggressively degraded in our growth conditions (Figure S5) (50,51). Hxt1 is a low-affinity hexose transporter expressed under high glucose conditions and is one of the major hexose transporters in *S. cerevisiae* (31). When expressed from a plasmid using the same expression cassette as *NC* and *OPT*, the growth rate for cells expressing *HXT1* and *HXT5* are quite different (Figure 5B). Cells expressing *HXT1* from the plasmid had similar growth rates as *OPT*, whereas cells expressing *HXT5* from the plasmid had higher growth rates similar to *NC*. Importantly, these growth assays were performed using yeast strain BY4741, not D452-2, indicating that the slow-growth phenotype for *OPT* is strain-independent. The growth rates for the endogenous transporters were correlated with activation of the UPR response, as observed for *NC* and *OPT* expression (Figure 5A). The protein concentration levels for Hxt1 and Hxt5, which were fused with eGFP at their C termini, were 2.3 and 1.6 times higher than the levels for Opt (Figure 5C). Thus, although some endogenous transporters can induce the effects seen with *OPT*, there does not seem to be a correlation between transporter protein levels and slower growth rates.

### The N-terminus of CDT-1 with optimized codons does not impact growth when fused to a cytoplasmic protein

We also investigated whether the slow-growth phenotype caused by the first 20-30 codons of *OPT* was an inherent property of this particular nucleotide sequence or whether the phenotype was due to other properties such as codon-usage in this region. We fused the first 30 codons of *NC* or *OPT* to the N-terminus of eGFP (*NC30-GFP* and *OPT30-GFP*) and expressed them with the same transcriptional regulation as *NC* and *OPT*. The growth rate of cells expressing these GFP fusions was identical (Figure 6A), despite the protein concentration for the Opt30-GFP fusion being 2.3-fold higher than Nc30-GFP and 7-fold higher than Opt (Figure 6B). These results indicate that the synonymous codon changes in *OPT* region 1 were only detrimental in the context of a membrane protein coding sequence.

**FIGURE 6.**
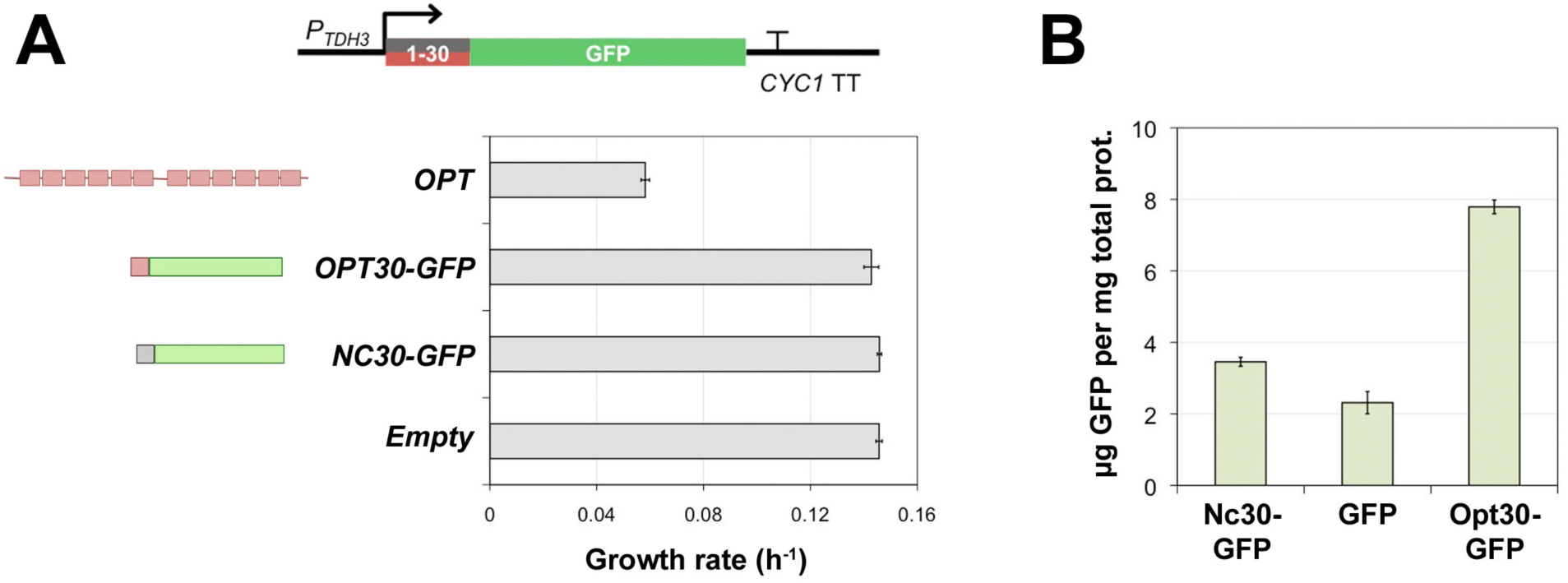
*NC30-GFP* and *OPT30-GFP* chimeras. (*A*) Growth rate of D452-2 cells expressing GFP chimeras. Region 1 (codons 1-30) from *NC* and *OPT* were fused to the N-terminus of GFP. *NC30-GFP* and *OPT30-GFP* were expressed from the same expression cassette as *NC* and *OPT*. Mean values of biological triplicates and standard errors are shown. (*B*) Protein levels in whole cell lysates for these chimeras were 3.45 ± 0.13 μg of GFP/mg total protein for Nc30-GFP, and 7.79 ± 0.19 μg of GFP/mg total protein for Opt30-GFP. Growth rates were calculated from biological triplicates.

### Differences in ribosome occupancy on NC and OPT transcripts

The *NC* and *OPT* sequences differ by only one nucleotide in the 5’-untranslated region (5’-UTR) of the mRNAs, having nearly identical Kozak and start codon contexts (5’ aacaaa<u>AUG</u>UCG/C for *NC/OPT*), suggesting that differences in translation initiation between mRNA versions are unlikely to cause the observed phenotypes (52). We therefore wondered whether changes in the elongation rate through the N-terminal region of the *OPT* sequence could affect subsequent steps in membrane protein biogenesis, since this region would encompass the slow translation ramp predicted to be a general feature of translation (17). Increases in translational speed through this region could interfere with productive protein synthesis, leading to proteostatic stress and decreased cell fitness. However, the slow translation ramp model is thought to affect the entire proteome, whereas we observe slow growth only when expressing *OPT* and not the *OPT30-GFP* chimera. To investigate whether codon usage in *OPT* might have led to an increase in elongation speed through the N-terminal region of CDT-1, we carried out ribosome profiling (RP) on D452-2 cells expressing *NC* or *OPT* (53,54), in order to make side-by-side comparisons of the ribosomal footprint distribution on *NC* and *OPT* mRNAs.

To enable analysis of the ribosomal footprint density at the 5’-end of the *CDT-1* open reading frame, we harvested yeast cells expressing *NC* or *OPT* at mid-exponential phase without the addition of cycloheximide to the cultures, and used rapid flash freezing (55). RP and mRNA libraries were prepared, sequenced and analyzed as previously described (Material and Methods; (53,54)). Information on replicate correlation can be found in Supplementary File 1. In the experiment, the mRNA levels of the *NC* and *OPT* versions of *cdt-1* were nearly identical (Figure 7A). We therefore normalized the ribosome footprint counts to reads per Million (rpM) and compared the read distribution throughout the protein (Figure 7B).

**FIGURE 7.**
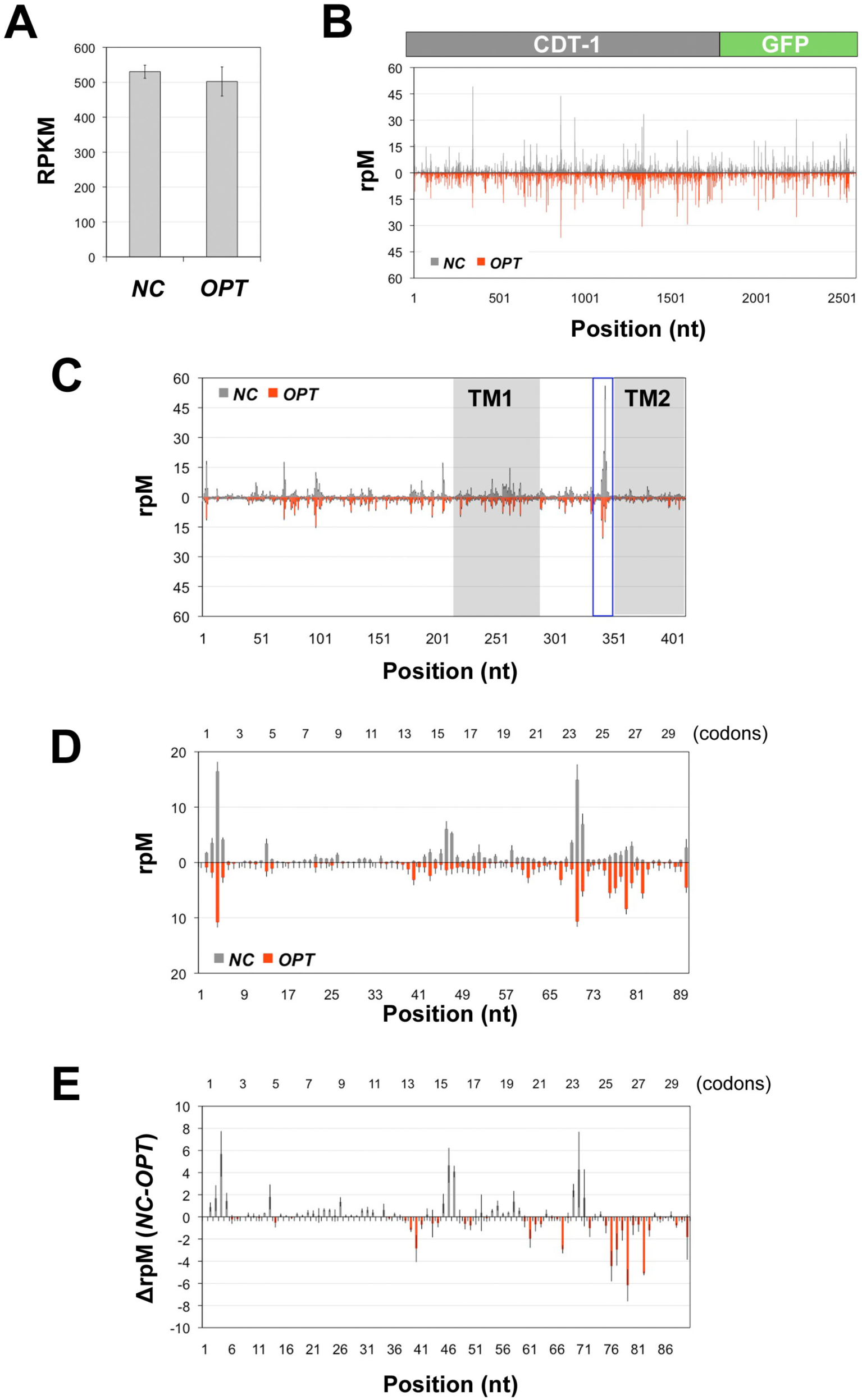
Ribosome footprint distributions for *NC* and *OPT* mRNAs. (*A*) mRNA RPKM levels for *NC* and *OPT* transcripts. The transcript levels for *NC* were 530.4 ± 18.9 RPKM; for *OPT* they were 502.3 ± 41.7 RPKM, determined from biological triplicates. (*B*) rpM counts at the A-site in the ribosome for both *NC* and *OPT* transcripts (these transcripts include the eGFP fusion). The x-axis represents the nt position in the coding sequence. (*C*) rpM counts for nucleotides 1-411 in *NC* and *OPT*. Shaded boxes represent the location of TM1 and TM2. The blue box marks the pause before TM2. (*D*) Ribosome footprint distribution for the first 90 nucleotides of the *NC* and *OPT* open reading frames. Shown are the mean rpM counts from biological triplicates with their corresponding standard errors. (*E*) Mean *OPT* rpM counts subtracted from the mean *NC* rpM counts (ΔrpM), and corrected for the standard error.

The most striking feature of the footprint distribution is the high footprint count at codons 113-114 in both *NC* and *OPT* transcripts (Figure 7C, blue box). This peak is 40-41 codons downstream of the predicted beginning of transmembrane helix 1 (TM1) in CDT-1 and is possibly an encoded REST element (16). However, the coding sequences around the pause region for both *NC* and *OPT* (^110^DTGPKVSV^117^) have high CAI values (0.761 for *NC* and 0.908 for *OPT*) (Figure S6), i.e. both regions are composed of preferred codons (19), whereas REST elements have been predicted to be encoded by less-preferred codons (16). A more likely explanation for the peak is SRP-induced elongation attenuation after binding TM1 (6,12,14). Interestingly, this peak is attenuated in the *OPT* sequence, which suggests that not all of the ribosome-nascent chain complexes were recognized by SRP.

### The Ribosome-Associated Complex interacts with Opt and triggers proteostatic stress

Since the *OPT* phenotype was only present in the context of membrane proteins, and did not affect cellular fitness during expression of cytosolic GFP chimeras, we wondered whether early membrane protein targeting interactions at the ribosome could be affected by altered elongation rates through the N-terminal sequence. We noticed that the ribosome footprint density for the first 72 nucleotides of the *OPT* open reading frame was much lower than that on *NC* through this region (Figure 7D, 7E, Figure S7). Since the mRNA levels were the same for both versions, the changes in footprint counts probably reflect faster translation through this region of the *OPT* transcript. For example, Yu and colleagues (2015) observed the same depletion of ribosome footprints with synthetic luciferase constructs that contained regions with the most commonly used codons from *N. crassa* (56).

Notably in CDT-1, the first 30 amino acids precede the signal sequence recognized by the SRP by more than one length of the ribosome exit tunnel (TM1 in CDT-1 begins at codon 73). We therefore focused our attention on complexes upstream of the SRP recognition step (Figure 8A). The nascent polypeptide-associated complex (NAC) modulates early interactions of SRP with nascent chains by preventing low affinity binding to non-secreted substrates and facilitating nascent chain sampling by SRP (9-11,57). NAC is a heterodimer of αNAC (encoded by *EGD2*) and βNAC (encoded by *EGD1* or *BTT1*) (58). Egd1 and Egd2 form the predominant dimer species of NAC and interact with ribosomes translating secreted and membrane proteins (11). Due to the increased elongation speed in the first 24 codons of *OPT* (Figure 7-figure supplement 2), we suspected that NAC could be mis-identifying the Opt polypeptide in the exit tunnel and blocking SRP from binding. To address this possibility, we expressed *NC* or *OPT* or an empty vector plasmid control in NAC knockout strains from the *Saccharomyces* Gene Deletion Project (*Δegd1* or *Δegd2*) (Figure 8A). We observed the same pattern of growth phenotypes on glucose as WT strains, with cells expressing the empty plasmid growing fastest, followed by cells expressing *NC*, and cells expressing *OPT* growing the slowest (Figure 8B). These results indicate that NAC is not involved in the slow growth phenotype induced by translation of the N-terminus of Opt.

**FIGURE 8.**
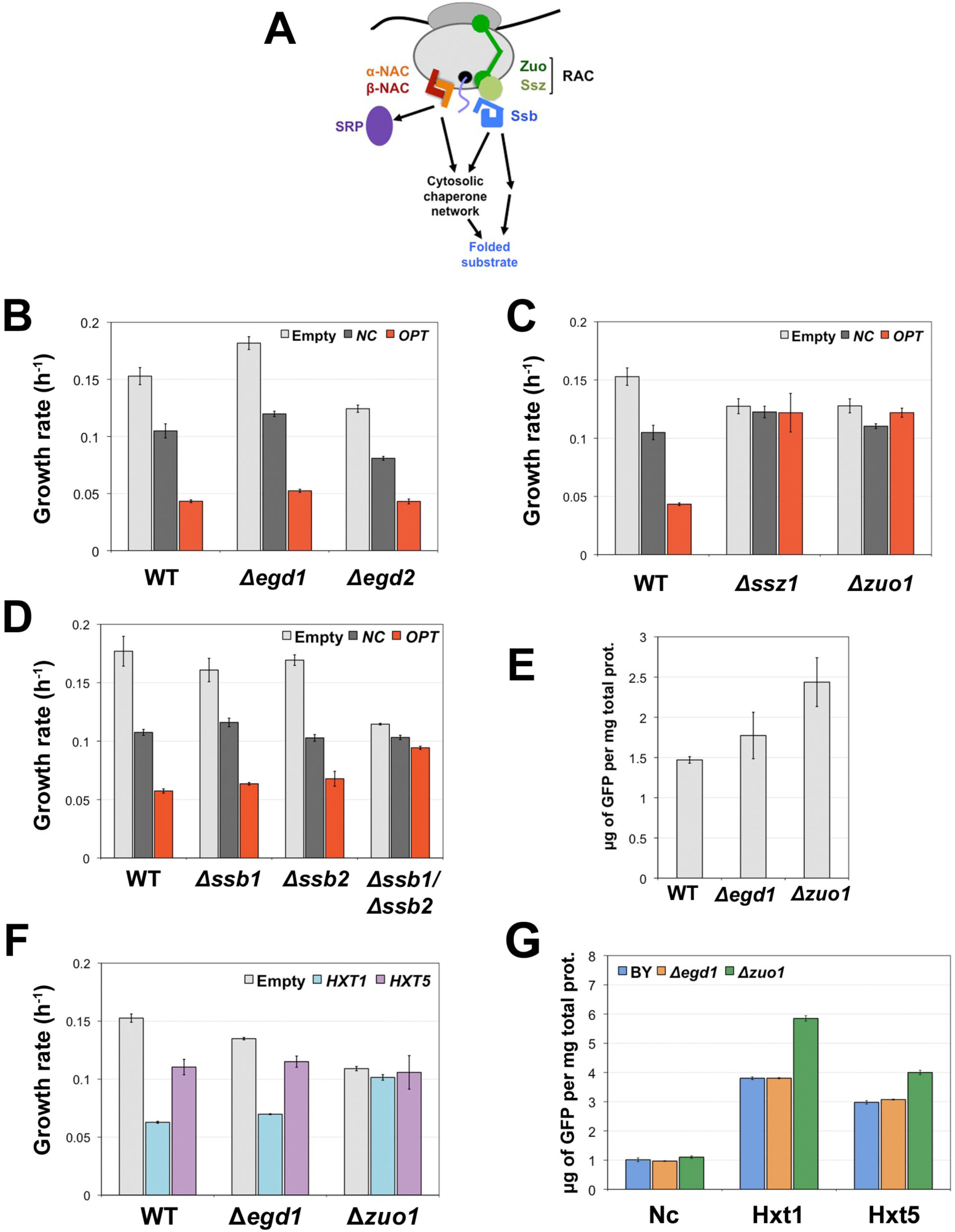
Contributions of chaperone activities to transporter expression and growth phenotypes. (*A*) Schematic of RAC and NAC interacting with the ribosome. NAC modulates SRP substrate recognition and chaperones a subset of nascent chains. RAC recruits Ssb to interact with nascent chain substrates and links aberrant proteins at the ribosome with the cytosolic network of chaperones (91). (*B*) Growth rates for WT, *Δegd1* and *Δegd2* strains expressing *NC* and *OPT.* (*C*) Growth rates for WT, *Δssz1* and *Δzuo1* strains expressing *NC* and *OPT.* (*D*) Growth rates for WT, *Δssb1, Δssb2*, and *Δssb1*/*Δssb2* expressing *NC* and *OPT*. (*E*) Opt protein concentration in WT, *Δegd1*, and *Δzuo1* whole cell lysates. Protein levels were quantified via GFP fluorescence. (*F*) Growth rates for WT, *Δegd1,* and *Δzuo1* strains expressing plasmid-borne *HXT1* and *HXT5*. All assays represent the mean and corresponding standard error from three biological replicates. (*G*) Nc, Hxt1, and Hxt5 protein concentrations in WT, *Δegd1*, and *Δzuo1* whole cell lysates. Protein levels were quantified via GFP fluorescence.

The other complex that is present at the ribosomal exit tunnel is the ribosome-associated complex (RAC). RAC and its cognate chaperones, the Hsp70 heat shock proteins Ssb1 and Ssb2 together form a chaperone triade and bind concurrently to the ribosome, but are thought to have a different subset of client nascent chains compared to NAC—mostly nuclear and cytoplasmic proteins (11,59), as well as ribosomal subunit assembly (60,61). RAC is formed by the Hsp70 homolog Ssz1 and the J Domain Hsp40 protein Zuo1 in *S. cerevisiae* (62,63). Surprisingly, although RAC has not been reported to interact with SRP substrates (59), *Δzuo1* and *Δssz1* cells (Figure 8A) expressing *NC* and *OPT* and grown on glucose had nearly identical growth rates and eliminated the slow-growth phenotype (Figure 8C). This phenotypic rescue is specific to expression of *NC* and *OPT*, since the *Δzuo1* and *Δssz1* cells expressing the empty plasmid control grew slightly slower than WT (Figure 8C). To confirm that this rescue was due to loss of the nascent peptide-Hsp70 chaperoning interaction, we assessed phenotypic changes in cells in which *ssb1* or *ssb2* or both were deleted (*Δssb1*, *Δssb2, Δssb1/Δssb2*), since RAC modulates cotranslational binding of Ssb1 and Ssb2 to client proteins (Figure 8A) (59). We measured growth rates in glucose for *Δssb1* and *Δssb2* cells expressing *NC* or *OPT*, and found that single deletion of either Hsp70 did not rescue the slow-growth phenotype in cells expressing *OPT* (Figure 8D). This was not unexpected as *SSB1* and *SSB2* are functionally redundant in yeast. However, deletion of both *SSB1* and *SSB2* (*Δssb1*/*Δssb2*), which had no impact on growth of *NC*-expressing cells, rescued the slow-growth defect in cells expressing *OPT* (Figure 8D). As with the *Δzuo1* and *Δssz1* cells, the rescue is specific to expression of OPT, since the *Δssb1*/*Δssb2* double deletion slowed growth of cells expressing the empty vector. The rescue observed in *Δzuo1*, *Δssz1* and *Δssb1*/*Δssb2* cells is also the opposite of that seen in RAC/Ssb deletion studies (64,65), including those which identified RAC/Ssb involvement in ribosomal subunit assembly (60,61). Taken together, these results indicate that recognition and binding of a subset of *OPT*-encoded proteins by the RAC-Ssb chaperone system led to proteostatic stress and decreased cell fitness. These results are consistent with the observed attenuation of the putative SRP pause in *OPT* transcripts after TM1 (Figure 7C).

To test whether RAC binding to a subset of Opt polypeptides possibly leads to their degradation, we measured Opt levels in WT, *Δegd1,* and *Δzuo1* cells. Notably, Opt levels were highest in lysates from the *Δzuo1* strain, while the *Δegd1* and WT Opt levels were similar (Figure 8E). These results suggest that RAC recognition targets a portion of the Opt polypeptides for degradation, as deletion of one of the components of RAC increased productive protein levels, and the C-terminal GFP fused to Opt would only be made after translation of Opt. As noted previously, over-expression of *HXT1* also led to a slow-growth phenotype and UPR activation, while *HXT5* expression under similar conditions did not (Figure 5). To test whether RAC may also interact with Hxt1, we transformed WT, *Δegd1*, and *Δzuo1* cells with pRS316-*HXT1*, pRS316-*HXT5*, or an empty plasmid as control, and measured their growth rates in glucose. Whereas the growth rate remained the same for all strains expressing *HXT5* from the plasmid, WT and *Δegd1* cells expressing *HXT1* from the plasmid had much lower growth rates (Figure 8F). Notably, as observed in *Δzuo1* cells expressing *OPT*, *Δzuo1* cells expressing *HXT1* from the plasmid grew at nearly the rate of cells expressing *HXT5* from the plasmid (Figure 8F). Consistent with the observations with *OPT* expressing cells, the phenotypic rescue in *Δzuo1* cells expressing *HXT1* is specific to expression of the transporter, as *Δzuo1* cells expressing the empty plasmid exhibited slower growth. Furthermore, Hxt1 and Hxt5 levels were higher in the *Δzuo1* strain and were similar in WT and *Δegd1*, while Nc levels remained the same in all three strains (Figure 8G). RAC therefore interacts with Hxt1 in the same manner as it does with Opt ribosome nascent-chain complexes.

### Ssb1 interacts with MFS transporter nascent chains at the ribosome before the emergence of TM1

We performed Ssb1-mediated selective ribosome profiling to identify substrates and analyze how RAC-Ssb interacts with nascent chains at the ribosome (66). For this analysis, we compared ribosome profiling data from two samples generated from the same culture: the translatome set, which includes footprints for all ribosome-nascent chain complexes; and the Ssb1 interactome set, which includes footprints from RNCs interacting with Ssb1 isolated after immunoprecipitation. We focused on the mRNAs for endogenous MFS transporters and were able to reliably identify eight transporters that interacted with Ssb1 during their translation (Figure 9; Figure S8; Table S4). Figure 9 shows the enrichment of *HXT1* footprints in interactome samples versus the translatome samples. There are two regions that are enriched early in translation where Ssb1 would interact with the nascent polypeptide at codon positions 74 and 93, before TM1 exits the tunnel. These results indicate that Ssb1 interacted with nascent chains of Hxt1 before the substrate binding site of SRP emerged (TM1). Further, nascent chains from the other 7 transporters that were identified (Arn3/Sit1, Arn4/Enb1, Hxt2, Hxt3, Itr1, Pho84, Tna1; Figure S8) had similar interactions with Ssb1: the chaperone bound very early in translation before or as TM1 was emerging from the peptide tunnel. We tried to survey other MFS transporters for interactions with Ssb1 (Table S5), but due to their inherent low expression we could not obtain a reliable signal for these transcripts.

**FIGURE 9.**
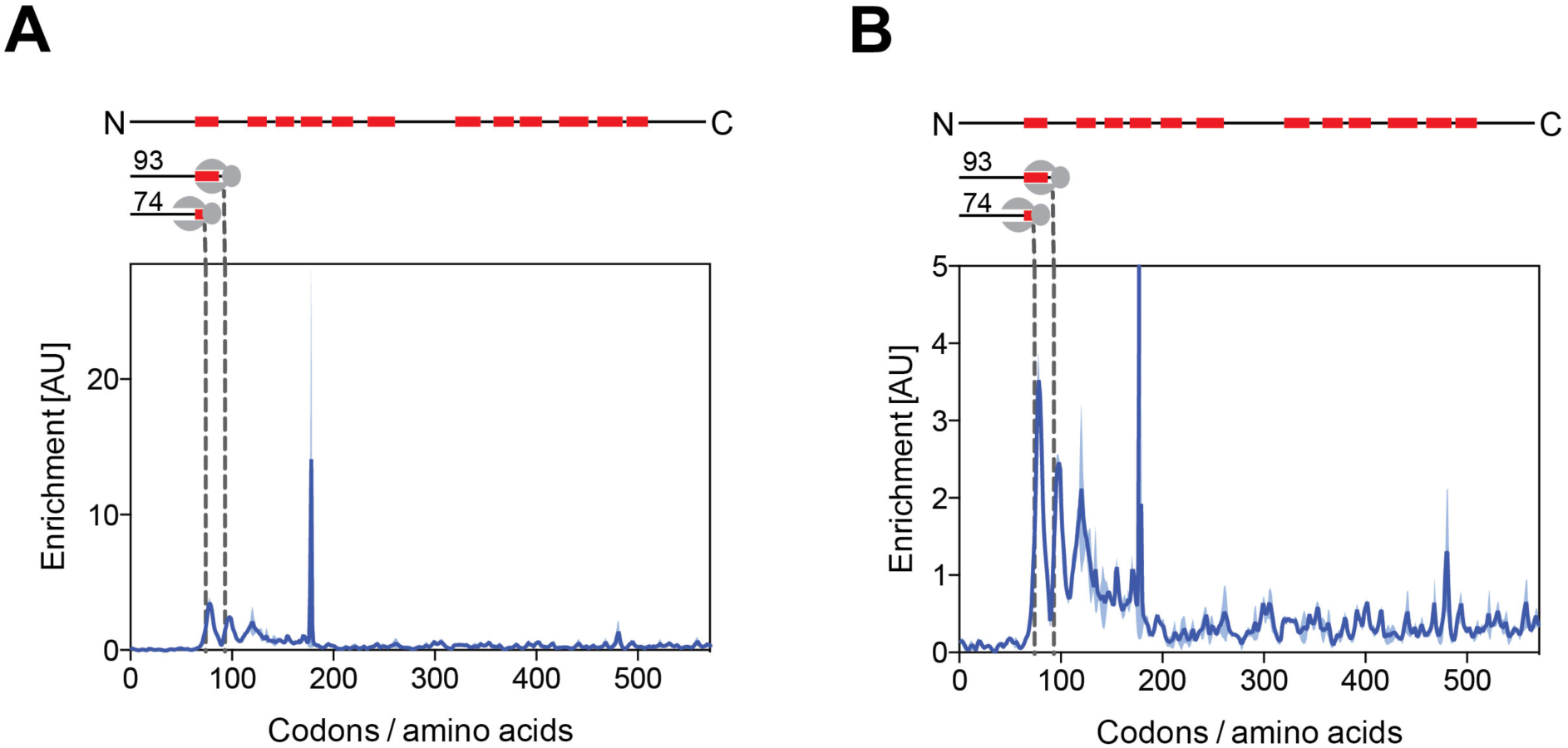
Ribosome footprint enrichment for *HXT1* from Ssb1-mediated selective ribosome profiling. (*A*) Footprint distribution over the entire *HXT1* transcript. Plot shows the enrichment per codon position in footprint counts in the interactome samples over the total translatome samples for *HXT1.* The top schematic shows primary structure of Hxt1 with the predicted positions for the transmembrane domains (TMDs, red boxes). Bottom schematic shows the length of the nascent chain in the ribosome at the peaks denoted by dashed lines. (*B*) Same plot zoomed in. Light blue shading indicates variance between biological duplicates.

## DISCUSSION

Although ER associated protein degradation (ERAD) serves as a quality control mechanism for membrane proteins post-translationally, less is known about quality control of membrane proteins before they are targeted to the translocon in the ER (25). Our results suggest that RAC-Ssb acts in the quality control of integral membrane proteins by actively sensing translation elongation rates of nascent membrane proteins on the ribosome. Indeed, cryo-EM reconstructions of RAC-80S ribosome complexes show that Zuo1 bridges the two subunits and extends from the ribosome peptide tunnel exit to the 40S subunit, where it interacts with the ES12 RNA helix (67).

RAC and Ssb are far more abundant than SRP in yeast cells, although both complexes interact dynamically with the ribosome (45,68). For RAC and Ssb, there are approximately 27 and 89 complexes per 100 ribosomes in exponentially growing *S. cerevisiae* cells, respectively. By contrast, there are only about 1-3 SRP complexes per 100 ribosomes under the same conditions (68). Furthermore, the large hydrophobic sequences in the first transmembrane helix (TM1) would present ideal binding regions for Ssb (59). Thus, the RAC-Ssb complex could easily outcompete SRP for binding a subset of Opt and Hxt1 nascent polypeptides. Consistent with this model, we observed binding of Ssb1 to nascent Hxt1 and 7 other MFS transporters polypeptides at the ribosome very early in translation (before or as TM1 emerges from the exit tunnel; Figure 9, Figure S8). Our results indicate that, for some integral membrane protein substrates, RAC-Ssb may compete with SRP at the ribosome. Indeed, in cryo-EM reconstructions of SRP-80S and RAC-80S complexes, SRP and RAC bind to proximal regions of the 60S subunit, which would prevent their simultaneous binding to the ribosome (67).

Several studies have shown that SRP’s interactions with substrates and the SRP receptor are kinetically constrained (7,45,46,69,70). For example, SRP interactions with substrates containing a signal sequence fall off dramatically after 140 amino acids have been translated (70). Further, SRP targeting to substrates with weak signal sequences is sensitive to translation elongation rates (69). In the context of integral membrane protein biogenesis, translation elongation rates may affect the sensitivity of SRP binding to the first transmembrane segment, TM1. Taking into account the substoichiometric cellular concentration of SRP compared to ribosomes, higher translation elongation rates prior to TM1 emergence may favor RAC-Ssb binding and preclude later SRP interrogation of the emerging nascent polypeptide. Our results suggest that another function of the ubiquitous slow translation ramp in the first 30-50 codons of membrane proteins (17) would be to minimize RAC recognition of membrane proteins and subsequent sorting to Ssb. Although we did not measure translation elongation rates on *HXT1* and *HXT5* mRNAs, we predict that the N-terminal codon usage in HXT1 has not been under evolutionary pressure to reduce translation speed. It will be interesting in the future to determine whether there is a negative correlation between endogenous MFS transporter expression level and translation speeds through the N-terminal sequence.

RAC-Ssb-bound Opt and Hxt1 nascent chains are probably targeted for degradation cotranslationally, as we observe increases in Opt and Hxt1 levels in *Δzuo1* strains (Figure 8E, 8G). Wang *et al.* observed cotranslational ubiquitination of nascent chains in actively translating ribosomes in human cells; however, the mechanism whereby these ubiquitinated nascent chains are delivered to the proteasome has not been elucidated to date (71). Our results suggest the subset of Opt nascent chains targeted by RAC are likely to be cotranslationally ubiquitinated, since we observed an increase in the concentration of Opt transporters in strains where a component of RAC (*Δzuo1*) was deleted (Figure 8E). Furthermore, systems-level analysis of *S. cerevisiae* nascent proteins targeted for cotranslational proteasomal degradation identified that these poly-ubiquitinated nascent chains generally had higher mRNA abundance and tAI— predictive of higher translation elongation rates (56,72)—than nascent polypeptides that were less susceptible to cotranslational ubiquitination (73). The cell may therefore use a quality control mechanism present at the ribosome to distinguish nascent polypeptides headed towards non-native folding species even after chaperone intervention. Our results suggest that RAC-Ssb may be responsible for this role in the context of at least MFS class of membrane proteins. When rapidly translated membrane proteins such as Opt are highly expressed, RAC-Ssb interactions with these nascent chains and the subsequent degradation of a subset of these polypeptides could overwhelm the basal level of protein quality control in cells, which would increase the proteasomal load and trigger cell stress and possibly the UPR (44,71). Notably, transporters were highly enriched in the subset of proteins identified to be toxic in a genome-wide analysis of protein overexpression in yeast (74), consistent with our model for RAC-Ssb function.

Our observations reveal another layer of complexity to expressing membrane proteins in heterologous systems in order to engineer novel metabolic pathways, especially when the host organism and the donor organism are phylogenetically distant. Even under autologous expression conditions, codon-optimization of the human wild-type cystic fibrosis transmembrane conductance regulator resulted in lower protein yields (75). Current theoretical metrics that rely on codon and tRNA frequencies (CAI and tAI) may not effectively predict whether a codon-optimized membrane protein sequence will have higher translational elongation rates. Pechmann and colleagues proposed a more comprehensive metric, which they termed the normalized translation efficiency scale (nTE) that takes into account the tAI for a specific sequence and the composition of the supply pool of tRNAs in the cell under a given growth condition (76). This scale may present a better method to predict translational efficiency under non-stressed conditions, and indeed, in a subsequent study, the authors showed that regions of low nTE after signal sequences in secreted proteins correlated with higher densities of ribosomal footprints (16). Nevertheless, under the stress conditions we observed in our experiments, the tRNA pool may vary drastically from normal conditions and these changes must be taken into account to increase the predictive power of the nTE scale under stress conditions (77,78). Previous reports have highlighted how silent polymorphisms in membrane protein coding sequences linked to human disease affect protein folding and function (22,23,79). Our observations provide a connection between these observations, translational efficiency and RAC-mediated quality control. The role of RAC-Ssb in membrane protein biogenesis also is likely to impact membrane protein expression in biotechnological applications.

## EXPERIMENTAL PROCEDURES

### Strains, plasmids and media

The *N. crassa* and the codon-optimized versions of *cdt-1*, and the *HXT* transporters were cloned into the pRS316 plasmid (*CEN URA*) using the In-Fusion HD Cloning Kit (Clontech Laboratories, Inc., Mountain View, CA). These vectors are pRS316-*NC*, -*OPT*, -*HXT1 or -HXT5*, respectively. The *NC* version of *cdt-1* was PCR-amplified from cDNA synthesized from mRNA isolated from *N. crassa* (FGSC 2489) grown on minimal media plus Avicel (microcrystalline cellulose) as the sole carbon source (36). Lifetech Geneart (ThermoFisher) performed gene synthesis and codon-optimization for the optimized version of *cdt-1*. *S. cerevisiae HXT* transporters were cloned from gDNA isolated from strain D452-2.

For all chimeras described, the region to be swapped was PCR-amplified from the donor sequence and cloned into a plasmid containing the flanking sequences from the recipient gene using the In-Fusion HD cloning kit. All genes were expressed under the control of *S. cerevisiae TDH3* promoter and the *CYC1* terminator; all MFS transporters were fused with eGFP at the C-terminus. Chimera 1 consisted of *NC* codons 1-170 and OPT codons 171-579; chimera 2 consisted of *OPT* codons 1-170 and 329-579, and *NC* codons 171-328; chimera 3 consisted of *OPT* codons 1-328 and 447-579, and *NC* codons 329-446; chimera 4 consisted of *OPT* codons 1-446, and *NC* codons 447-579.

*S. cerevisiae* strains BY4741, BY4741 deletion strains [from the *Saccharomyces* Genome Deletion Project at Stanford University (80)], D452-2 (81), and D452-2 UPR reporter were used in this study. BY4741 *Δssb1*, *Δssb2*, and *Δssb1Δssb2* were constructed for this study. In the case of the *Δssb1* and *Δssb2* strains, deletion of the respective gene was carried out following the same deletion design used to create the *Saccharomyces* Genome Deletion library. Briefly, the *KanMX* cassette was cloned from the gDNA of the *Δzuo1* strain with primers that had 50-bp homology to the promoter and terminator regions of *SSB1* and *SSB2,* and transformed into WT BY4741 cells. *Δssb1* was then used as the parental strain to create the *Δssb1Δssb2* deletion strain; *SSB2* was deleted via CRISPR-Cas9 using a protocol developed by (37). The D452-UPR reporter strain was constructed as follows. The UPR reporter was cloned from genomic DNA from strain YJW1200 provided by Jonathan Weissman’s lab at UCSF (42,82). Details of reporter construction are provided in (42). The *4XUPRe-GFP*-*TEF2pr*-*RFP* reporter cassette was inserted into the *HO1* endonuclease locus of D452-2 using CRISPR-Cas9 (37). A list of primers for cloning and Cas9 guides is provided in Supplemental File 2, Table 2. Yeast cells were transformed using the Frozen-EZ Yeast transformation kit (Zymo Research Corp., Irvine, CA, USA).

For yeast growth experiments, optimized Minimal Media (oMM) lacking uracil (40) was used. oMM contained 10 g/L (NH4)_2_SO_4_, 1 g/L MgSO_4_ · 7H_2_O, 6 g/L KH_2_PO_4_, 100 mg/L adenine hemisulfate, 30 g/L glucose or 20 g/L cellobiose, 1.7 g/L YNB (Sigma-Aldrich, Y1251), 2x recommended CSM-Ura dropout mix (MP Biomedicals, Santa Ana, CA), 10 mg/L of inositol, 100 mg/L glutamic acid, 20 mg/L lysine, 375 mg/L serine, 100 mM morpholineethanesulfonic acid (MES), pH 5.7.

### Codon usage analysis

Codon usage tables for *N. crassa* and *S. cerevisiae* were obtained from the Codon Usage Database (http://www.kazusa.or.jp/codon/) (83). The *NC* sequence was analyzed using the %MinMax web-server (http://www.codons.org/) (38). CAI calculations were carried out using the CAIcal server (http://genomes.urv.es/CAIcal/) (39).

Per codon tAI was calculated from the nucleotide sequence using the tAI calculator (http://web.stanford.edu/group/frydman/codons/co dons.html) (76).

### Growth assays

All growth assays were performed using the Bioscreen C (Oy Growth Curves Ab Ltd., Helsinki, Finland) with biological triplicates or quadruplicates. Single colonies of *S. cerevisiae* strains transformed with pRS316 containing the MFS transporter of interest were grown in oMM-Ura to late-exponential phase at 30 °C in 24-well plates. Cultures were pelleted at 4000 rpm, the spent media supernatant was discarded and cells were resuspended in H_2_O. The OD was measured to calculate the inoculum volume needed for 200 μL cultures at an initial OD ≈ 0.1 in Bioscreen plates. The OD at 600 nm was measured in 10 min intervals for 48-72 h at 30 °C. Growth curves were fit and growth parameters were calculated using the ‘grofit’ package in R (84), (https://www.r-project.org/).

### Microscopy

D452-2 cells expressing *NC* or *OPT* were grown in oMM-Ura media to mid-exponential phase at 30 °C. The cultures were centrifuged, spotted onto glass slides and examined on a Leica DM 5000B Epi-fluorescence microscope at 100x DIC (Leica, Germany). Transporters were visualized using the L5 filter cube; images were captured using the Leica DFC 490 camera and analyzed with the accompanying microscope software.

### Yeast-cell based cellobiose uptake assay

Yeast strain D452-2 cells transformed with pRS316-*NC* or pRS316-*OPT* were grown to mid-exponential phase. Cells were harvested and washed twice with Transport Assay Buffer (5 mM MES, 100 mM NaCl, pH 6.0) and resuspended to a final OD_600_ of 40. 500 μL of cell resuspension was quickly mixed with equal volume of Transport Assay Buffer containing 400 μM cellobiose. For the initial time point (t = 0 sec), an aliquot was removed and centrifuged for 1 min at 4 °C at high speed to pellet the cells and the supernatant was removed. The remaining cell resuspension was incubated at 30 °C for 30 min with constant shaking. After incubation, samples were centrifuged for 5 min at 4 °C at 14000 rpm and supernatant was removed. For analysis, 400 μL of supernatant were mixed with 100 μL of 0.5 M NaOH, and cellobiose concentrations remaining in the supernatant were measured on a Dionex ICS-3000 HPLC (ThermoFisher) using a CarboPac PA200 analytical column (150 x 3 mm) and a CarboPac PA200 guard column (3 x 30 mm) at room temperature. 25 μL of sample was injected and run at 0.4 mL/min with a mobile phase using 0.1 M NaOH.

### Protein concentration determination using fluorescence emission spectroscopy

Fifty mL cultures were grown at 30 °C until they reached an OD_600_ of 3, at which point they were harvested by centrifugation, resuspended in ∼4-5 mL of media and aliquoted into microcentrifuge tubes to yield a total OD_600_ of 30. Samples were spun down at 14,000 rpm for 1 min and the supernatant was aspirated. Cell pellets were quickly flash frozen in liquid N_2_. Frozen cell pellets were thawed on ice and 400 μL of Buffer A (25 mM Hepes, 150 mM NaCl, 10 mM cellobiose, 5% glycerol, 1 mM EDTA, 0.2X HALT protease inhibitor cocktail (ThermoFisher, Waltham, MA), pH 7.5) were added for resuspension. Cells were lysed with zirconia/silica beads in a Mini-Beadbeater-96 (Biospec Products, Bartlesville, OK). Cell debris was pelleted at 10,000x*g* for 10 min at 4 °C, lysates were diluted three-fold with Buffer A, and their GFP fluorescence was measured using a Horiba Jobin Yvon Fluorolog fluorimeter (Horiba Scientific, Edison, NJ). The λ_EX_ was 485 nm, and the emission wavelength was recorded from 495-570 nm, with both excitation and emission slit widths set to 3 nm. A fluorescence calibration curve was prepared with eGFP purified from *E. coli* (>95% purity). The settings and the eGFP protein concentration range were chosen to yield a linear correlation between the fluorescence intensity at the maximum λ_EM_ (510 nm) and the protein concentration of the standard. The maximum fluorescence intensity of the samples fell within this range. Target protein concentrations represent the mean from 3 biological replicates. Total protein concentration of the lysate was determined using the Pierce BCA Protein Assay Kit (ThermoFisher).

### In-gel Fluorescence Detection

BY4741 cells expressing pRS316-*HXT1* and pRs316-*HXT5* were grown to OD_600_ of 3, pelleted and flash frozen in liquid N_2_. Cells were resuspended in 400 μL of Buffer A and lysed using zirconia/silica beads in a Mini-Beadbeater-96. Cell debris was pelleted at 10,000x*g* for 10 min at 4 °C, and total lysate protein concentration was measured using the Pierce BCA Protein Assay kit. Lysates were diluted to 2 mg/mL, 4X SDS-PAGE Loading buffer was added, and 30 μL of samples were loaded immediately after loading buffer addition without boiling into a 4-20% SDS-PAGE gel. A standard consisting of purified recombinant GFP was used as a size control. Gels were run at 100 V for ∼1.5 h. After electrophoresis, the gel was rinsed with nanopure H_2_O and imaged using an ImageQuant LAS-4000 Imager (GE Healthcare Life Sciences, Marlborough, MA). The imager was set to the SYBR-SAFE fluorescence setting, with the light source set to 460 nm Epi, the excitation filter set to 515 nm, and the iris set to F 0.85.

### Fluorescence size-exclusion chromatography (FSEC)

Three separate 0.8 L cultures of oMM-Ura were inoculated with 10 mL seed cultures of D452-2 cells transformed with either pRS316-*NC* or -*OPT*. These cultures were grown to late-exponential phase at 30 °C and harvested at 4000x*g* for 10 min at 4 °C. Cells were resuspended in 50 mL of Buffer A, flash frozen in liquid N_2_ and stored at -80 °C. Cells were thawed on ice and lysed using 0.5 mm zirconia/silica beads in a BeadBeater (Biospec Products) at 4 °C. Cell debris was pelleted at 6000x*g* and the supernatant was centrifuged at 125,000x*g* for 1 h at 4 °C. The supernatant was discarded and the membrane pellet was resuspended via dounce homogenization in Buffer A. Resuspended membranes were stored at -80 °C.

Hattori et al. developed and optimized the thermostability FSEC analysis for membrane proteins fused with a C-terminal GFP fusion (41). The protocol used here is similar, with the following modifications. Membrane protein fractions (1600 μL) were solubilized by adding 10% n-dodecyl-β-D-maltoside (DDM) to a final concentration of 2% and incubating at 4 °C for 1.5 h in a rotating mixer. Ninety μL were aliquoted into 15 thin-walled PCR tubes and samples were heated to the target temperature for 10 min in an Applied Biosystems Thermal Cycler (ThermoFisher) and then cooled down to 4 °C. The thermal cycler lid temperature was kept at 100 °C. Five aliquots were then pooled and centrifuged at 100,000x*g* for 30 min at 4 °C; 300 μL of clarified supernatant were collected (resulting in technical triplicates for each temperature). From the remaining non-heated sample, a 400 μL control sample incubated only at 4 °C was collected; this sample was also centrifuged at 100,000x*g* after solubilization.

For FSEC, a Superose 6 10/300 GL SEC column (GE Healthcare Life Sciences, Pittsburg, PA) was attached to an Agilent 1200 HPLC system with a Fluorescence Detector module (Agilent Technologies, Santa Clara, CA). GFP fluorescence chromatograms were recorded at an excitation wavelength λ_EX_ of 485 nm, an emission wavelength λ_EM_ of 512 nm, with the output set to 1 V and the PMT gain to 12. One-hundred μL fractions were loaded onto the Superose 6 column, the flow rate was 0.4 mL/min and the chromatography runs were carried out at 22 °C. Chromatograms were analyzed using the “ChemoStation for LC 3D Systems” software from Agilent Technologies. The peak area for the peak corresponding to CDT-1 was calculated; the fraction folded was calculated from the ratio of the peak area at the target temperature to the peak area of the corresponding control incubated at 4 °C. The FSEC-Tm values were determined as the temperature at which half the CDT-1 protein remained folded. All samples for a temperature set were run consecutively on the same day. Tm values were calculated via non-linear regression analysis after fitting the Boltzmann sigmoidal equation to the data.

### Flow cytometry and UPR reporter measurements

All UPR reporter experiments were carried out using the D452-UPR reporter strain as described previously (42) with minor modifications. The UPR reporter strain was transformed with a pRS316 plasmid carrying the specified protein with a C-terminal GFP tag (P_*THD3*_ and *CYC1* terminator). In these plasmids, an R97S mutation was introduced into eGFP to eliminate fluorescence (85). We confirmed that this mutation abolished GFP fluorescence by expressing transporter-(R97S)eGFP in D452-2 cells and then carrying out flow cytometry measurements.

Transformed colonies were grown overnight in oMM-Ura in 24-well plates at 30 °C. The following day, fresh oMM-Ura cultures were inoculated with these overnight seeds and grown at 30 °C to a final OD_600_ of 3-4 (mid-exponential phase). Cultures were then centrifuged at 2500x*g*, media was discarded and cells were resuspended in 1X PBS buffer. Flow cytometry measurements were carried out subsequently and samples were analyzed in a BD Bioscience LSR Fortessa X20 (Flow Cytometry Facility at UC Berkeley). The number of events was stopped at 20,000 counts. Data were collected using FITC parameter settings for GFP (488 nm laser, 505 nm LP filter) and PE-Texas Red for RFP (561 nm laser, 595 nm LP filter). FCS files were converted to CSV files using the ‘FcstoCsv’ module in the GenePattern Server from the Broad Institute at MIT (86). These files provided the raw data for further analysis, which consisted of the pulse heights in the GFP and RFP channels for each event. The median for the log_2_ ratio of GFP/RFP for all events in a sample was used as a proxy for UPR reporter activation levels. All experiments were carried out using four or five biological replicates.

### Cell harvesting for ribosome profiling and RNA sequencing

For both ribosome profiling and RNA sequencing procedures, the same cell cultures were used. Five hundred mL cultures of oMM-Ura were inoculated with D452-2 cells transformed with pRS316-*NC* or -*OPT* at 30 °C. Biological triplicates were concurrently grown for each condition (*NC* or *OPT*). Cultures were harvested when they reached an OD_600_ of 3 via filtration using a Millipore filter device and a 0.8 μm filter (GE Healthcare Life Sciences). Cell patties were quickly scraped off the filter and submerged in liquid N_2_; no cycloheximide was added to the cultures prior to harvesting. Polysome lysis buffer (20 mM Tris, 140 mM KCl, 1.5 mM MgCl_2_, 200 μg cycloheximide, 1% Triton X, pH 8) was slowly dripped into liquid N_2_ tubes containing cells. Samples were stored at -80 °C until future use. Cycloheximide was added to the lysis buffer, and a portion of harvested cells was used for total mRNA sequencing.

### RNA Sequencing

The RiboPure Yeast kit (Ambion, Austin, TX) was used to extract total RNA from 40 OD of cells following the manufacturer’s instructions. Four μg of total RNA were used to prepare multiplexed libraries with barcodes using the TruSeq RNA Sample Prep Kit (Illumina). The final cDNA libraries were quantified using an Agilent Bioanalyzer 2000 (Fuctional Genomics Laboratory, UC Berkeley) and sequenced with the Illumina Genome Analyzer-II using standard Illumina operating procedures (Vincent J. Coates Genomic Sequencing Laboratory, UC Berkeley). Sequence reads were assembled and analyzed in CLC Genomics Workbench 6.5 (CLC Bio, Aarhus, Denmark). The genome for *S. cerevisiae* S288C (version R64.2.1), the parent strain of D452-2, was downloaded from Refseq at the NCBI (http://www.ncbi.nlm.nih.gov/refseq/); the mitochondrial genome was included. The sequence for pRS316-*NC* and -*OPT* were manually annotated and added to this file. Expression values were normalized by calculating the reads per kb of mRNA per million mapped reads (RPKM), and normalized further by using the option of “By totals”.

### Ribosome profiling

Preparation of ribosome profiling libraries followed similar protocols as those in (53,54). Frozen cell pellets were lysed by low-frequency cryogenic mixer milling. Lysate was clarified of cell debris and 25 A_260_ units of extract were treated with 450 U of RNase I (Ambion) for 1 h at room temperature with gentle rotation; digestion was stopped by addition of 120 U of Superase-IN. Ribosomes were pelleted using high-speed centrifugation through a 1 M sucrose cushion. miRNeasy Mini kit (Qiagen) was used to purifiy ribosome-protected mRNA fragments following the manufacturer’s instructions. After size selection and dephosphorylation, a Universal miRNA cloning linker (New England Biolabs) was ligated to the 3’ end of footprints, followed by reverse transcription, and circular ligation. Subtraction of rRNA fragments was performed using anti-sense biotinylated oligos for the most common rRNA fragment contaminants. After rRNA subtraction, the cDNA library was PCR-amplified. The final DNA libraries from biological triplicates for each condition (*NC* and *OPT*) were quantified using an Agilent Bioanalyzer 2000 (Fuctional Genomics Laboratory, UC Berkeley) and sequenced with the Illumina Genome Analyzer-II using standard Illumina operating procedures (Vincent J. Coates Genomic Sequencing Laboratory, UC Berkeley).

For footprint analysis, trimmed sequencing reads without the linker were aligned to the *S. cerevisiae* ribosomal sequences using Bowtie (87). These reads were removed, and the non-rRNA reads were then mapped to the *S. cerevisiae* genome using Tophat (88). The sequence for the pRS316-*NC/OPT* plasmids were manually annotated and added to separate genome fasta files created for each condition and designated as new chromosomes. Only uniquely aligned reads were used for subsequent analyses. Most of the reads were between 27-32 nt long and these reads were mapped onto their respective coding sequences as described previously (53,54). For rpM normalization, reads were normalized to the total number of reads mapped. For ΔrpM, the mean rpM for each position in *OPT* was subtracted from the mean rpM for the corresponding position in *NC*.

### SSB-mediated selective ribosome profiling

SeRP library preparation and analysis are described in detail in (66). The total mapped reads per library were:

translatome 1: 10,504,232

translatome 2: 9,249,833

interactome 1: 13,161,962

interactome 2: 20,801,425

Genes with fewer than 64 raw reads in the translatome or interactome were excluded from the analysis. MFS transporters in the yeast genome are included in Table S4 and Table S5. Ratios of RPM-normalized interactome and translatome data were built over a window of 15 nucleotides for both replicates. The average of both replicates was plotted and the variance between replicates was indicated using shading. The locations of transmembrane helices were predicted using MFS primary sequences and structure prediction using Phyre2 (89).

## Acknowledgement

We thank Gloria Brar, Ze Cheng, Nicholas McGlincy and Liana Lareau for helpful discussions, Jeremy Thorner for strains from the *Saccharomyces* Genome Deletion library, Jonathan Weissman for the UPR reporter strains, Hector Nolla for help with flow cytometry experiments and Stefan Bauer and Ana Ibañez Zamora for help with analytical methods. This work was supported by funding from the Energy Biosciences Institute (J.H.D.C.), and from the NIH (R01-GM065050)(J.H.D.C) and research grants from the Deutsche Forschungsgemeinschaft (SFB638 and FOR1805) (G.K. and B.B.).

## Competing interests

The authors declare that they have no conflicts of interest with the contents of this article.

## Author contributions

LAS conducted most of the experiments, analyzed the results, and wrote most of the paper. LAS and JHDC conceived of the project and experimental plans. KD, BK, and GC conceived, conducted experiments, and analyzed results dealing with SSB-mediated selective ribosome profiling. YL conducted RNA sequencing experiments and analyzed data. VYY conducted experiments dealing with promoter strength and protein expression.

